# Miro-dependent mitochondrial pool of CENP-F and its farnesylated C-terminal domain are dispensable for normal development in mice

**DOI:** 10.1101/415315

**Authors:** Martin Peterka, Benoît Kornmann

## Abstract

CENP-F is a large, microtubule-binding protein that regulates multiple cellular processes including chromosome segregation and mitochondrial trafficking at cytokinesis. This multiplicity of function is mediated through the binding of various partners, like Bub1 at the kinetochore and Miro at mitochondria. Due to the multifunctionality of CENP-F, the cellular phenotypes observed upon its depletion are difficult to interpret and there is a need to genetically separate its different functions by preventing binding to selected partners. Here we engineer a CENP-F point-mutant that is deficient in Miro binding and thus is unable to localize to mitochondria, but retains other localizations. We introduced this mutation in cultured human cells using CRISPR/Cas9 and show it causes a defect in mitochondrial spreading similar to that observed upon Miro depletion. We further create a mouse model carrying this CENP-F variant, as well as truncated CENP-F mutants lacking the farnesylated C-terminus of the protein. Importantly, one of these truncations leads to ∼80% downregulation of CENP-F expression. We observe that, despite the phenotypes apparent in cultured cells, mutant mice develop normally. Taken together, these mice will serve as important models to study CENP-F biology at organismal level. In addition, because truncations of CENP-F in humans cause a lethal disease termed Strømme syndrome and because CENP-F is involved in cancer development, they might also be relevant disease models.

## Introduction

CENP-F is a large coiled coil protein that was originally found as a kinetochore binding protein (Rattner et al., 1993). It is apparent from recent work that CENP-F also functions in mitochondrial transport, nuclear envelope breakdown, microtubule polymerization, and transcriptional regulation via its interactions with the retinoblastoma protein and ATF4 (Feng et al., 2006; Kanfer et al., 2015; Ma et al., 2006; Varis et al., 2006). Despite its multitude of subcellular localizations and binding partners, the physiological function of CENP-F remains poorly defined. To our knowledge, the only known phenotype observed upon CENP-F loss is dilated cardiomyopathy described in a heart-specific conditional knock-out mice (Dees et al., 2012).

In humans, aberrant expression of CENP-F has been implicated in prostate cancer (Aytes et al., 2014). In addition, mutations in CENP-F are known to cause Strømme syndrome, a rare autosomal recessive disorder characterized by microcephaly, intestinal atresia and other ciliopathy phenotypes (Filges et al., 2016; Ozkinay et al., 2017; Waters et al., 2015).

Both the expression level and subcellular localization of CENP-F are regulated in a cell cycle-dependent manner. Undetectable in G1, CENP-F accumulates in the nucleus during S/G2. At this stage, a fraction of CENP-F is exported and recruited to the outer nuclear envelope (NE) where it interacts with the nucleoporin Nup133 and participates in the recruitment of dynein (Bolhy et al., 2011). Subsequently in prometaphase, CENP-F relocates to kinetochores (Rattner et al., 1993). The molecular mechanism of CENP-F kinetochore binding has been studied in great detail (Berto et al., 2018; Ciossani et al., 2018; Zhu, 1999). CENP-F is targeted to the outer kinetochore via a KT-core domain (residues 2792 to 2887, Fig 1A) via synergistic action of Bub1 kinase and the kinesin Cenp-E (Berto et al., 2018; Zhu, 1999). In turn, CENP-F recruits Ndel1/Nde1/Lis1/Dynein complex necessary for chromosome segregation. In addition, a recent report showed that CENP-F is a receptor for ATR at the kinetochore and therefore an upstream trigger of the ATR-Chk1-Aurora B pathway necessary for the proper segregation of chromosomes (Kabeche et al., 2017). Nevertheless, the relevance of CENP-F at the kinetochore is controversial. Numerous studies reported defects in chromosome segregation and/or cell cycle progression upon depletion of CENP-F or inhibition of its kinetochore localization (Bomont et al., 2005; Evans et al., 2007; Feng et al., 2006; Holt et al., 2005; Laoukili et al., 2005; Vergnolle and Taylor, 2007; Yang et al., 2005). While conflicting reports showed no defects in mitosis upon CENP-F loss (Ciossani et al., 2018; McKinley and Cheeseman, 2017; Pfaltzgraff et al., 2016; Raaijmakers et al., 2018).

**Figure 1:**
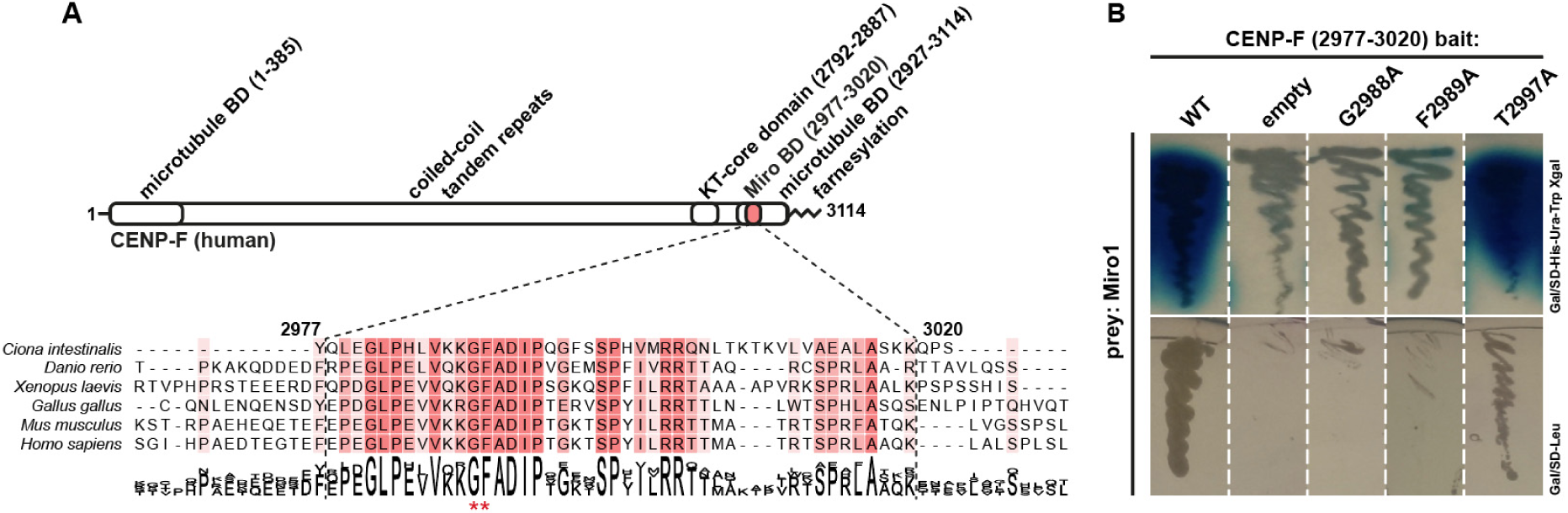
Disruption of CENP-F-Miro complex by single point mutations in CENP-F (**A**) Domain organization of CENP-F and alignment of Miro-binding domains from chordates. Red asterisks mark residues G2988A and F2989A essential for Miro binding. (**B**) Yeast two-hybrid assay using strains containing denoted plasmids. Top row: X-gal overlay assay, bottom row: growth assay on leucine dropout medium.

After exerting its function at kinetochores in mitosis, CENP-F is recruited in cytokinesis to the outer mitochondrial membrane by the atypical GTPases Miro1 and Miro2 (Kanfer et al., 2015). The Miro-binding domain is highly conserved and located near the C-terminus (2977-3020, Figure 1A) of CENP-F. Mechanistically, CENP-F appears to be linking mitochondria with the growing microtubule tips and harnessing the force generated by microtubule growth (Kanfer et al., 2015; Kanfer et al., 2017). This process helps the mitochondrial network to properly distribute throughout the cytoplasm during cytokinesis. Upon CENP-F depletion, mitochondria fail to spread to the cell periphery and remain clumped in the perinuclear area, phenocopying Miro depletion. Of note, a fraction of CENP-F is localized on mitochondria also in G2 (Kanfer et al., 2015; Kanfer et al., 2017). The physiological function of the mitochondrial fraction of CENP-F remains unknown.

Bracketing its coiled-coils, CENP-F harbors two microtubule-binding domains of unknown functions at both termini of the protein (Feng et al., 2006). Both domains have microtubule tip-tracking properties and a unique ability to transport cargo *in vitro* continuously with both growing and shrinking microtubules, with the C-terminus being a more robust mediator of these movements (Kanfer et al., 2017; Volkov et al., 2015). In cells, full-length CENP-F can track growing microtubule tips (Kanfer et al., 2017). These properties of CENP-F further substantiate its potential role in supporting microtubule tip-mediated transport of cellular cargoes such as chromosomes and mitochondria.

Additionally, CENP-F contains a CaaX motif, which is farnesylated (Ashar et al., 2000). However, functional consequences of CENP-F farnesylation have been debated. The CaaX motif of CENP-F has been proposed to be necessary for its kinetochore and NE localization (Hussein and Taylor, 2002; Schafer-Hales et al., 2007). Moreover, inhibition of CENP-F farnesylation led to defective degradation of the protein and delayed G2/M progression in cancer cells (Gurden et al., 2010; Hussein and Taylor, 2002). On the other hand, more recent studies showed only a mild or no effect of farnesylation on CENP-F kinetochore localization (Holland et al., 2015; Moudgil et al., 2015). Thus, the function and physiological relevance of CENP-F farnesylation remain unresolved.

The plethora of interacting partners and cellular localizations make studying CENP-F a challenge, and the cellular phenotypes observed upon CENP-F depletion are difficult to interpret due to the multifunctionality of the protein. Here, using a single point mutation within the Miro-binding region of CENP-F, we genetically separated the mitochondrial function from other functions of CENP-F. We used CRISPR/Cas9-mediated mutagenesis to introduce this mutation in human cells. Furthermore, to gain a physiological insight into mitochondrial function of CENP-F, we engineered a similar mutation in mice. Moreover, as a by-product of CRISPR/Cas9-mediated engineering, we generated animals bearing truncated CENP-F alleles lacking the last two exons of CENP-F, which encode the C-terminal microtubule-binding domain and the farnesylation motif. In addition to the loss of the C-terminal domain, one of the mutations resulted in approximately 80% decrease of overall CENP-F expression. Strikingly, despite the plethora of functions attributed to CENP-F, these mice are viable and fertile, and do not display any obvious phenotype. These different mouse models will be instrumental to systematically study the multiple functions of CENP-F in different tissues and under different physiological or pathological conditions.

## Results

We have previously mapped the Miro1/2-binding domain in CENP-F to a C-terminal region spanning residues 2977-3020 of human CENP-F, which is among the best conserved parts of the protein (Figure 1A) (Kanfer et al., 2015). Here, in order to specifically disrupt the interaction between CENP-F and Miro, we used alanine-scanning mutagenesis in the yeast two-hybrid assay and found that alanine substitution of the highly conserved CENP-F residue G2988 or F2989 completely abrogated Miro binding. On the other hand, alanine substitution of the non-conserved residue T2997 did not influence Miro1 binding in this assay (Figure 1B). Therefore, the interaction between CENP-F and Miro in yeast two-hybrid assay can be disrupted by a single point mutation, offering an opportunity to genetically separate the mitochondrial from other functions of this large multifaceted protein.

To characterize the properties of CENP-F^F2989A^ in human cells, we engineered the F2989A substitution into the endogenous *CENP-F* locus using CRISPR/Cas9-mediated genome editing (Figures 2A, B, C). We performed the substitution in human osteosarcoma cell line U2OS, which displays high levels of Miro-dependent mitochondrial recruitment of CENP-F in S/G2 and cytokinesis (Kanfer et al., 2015).

**Figure 2:**
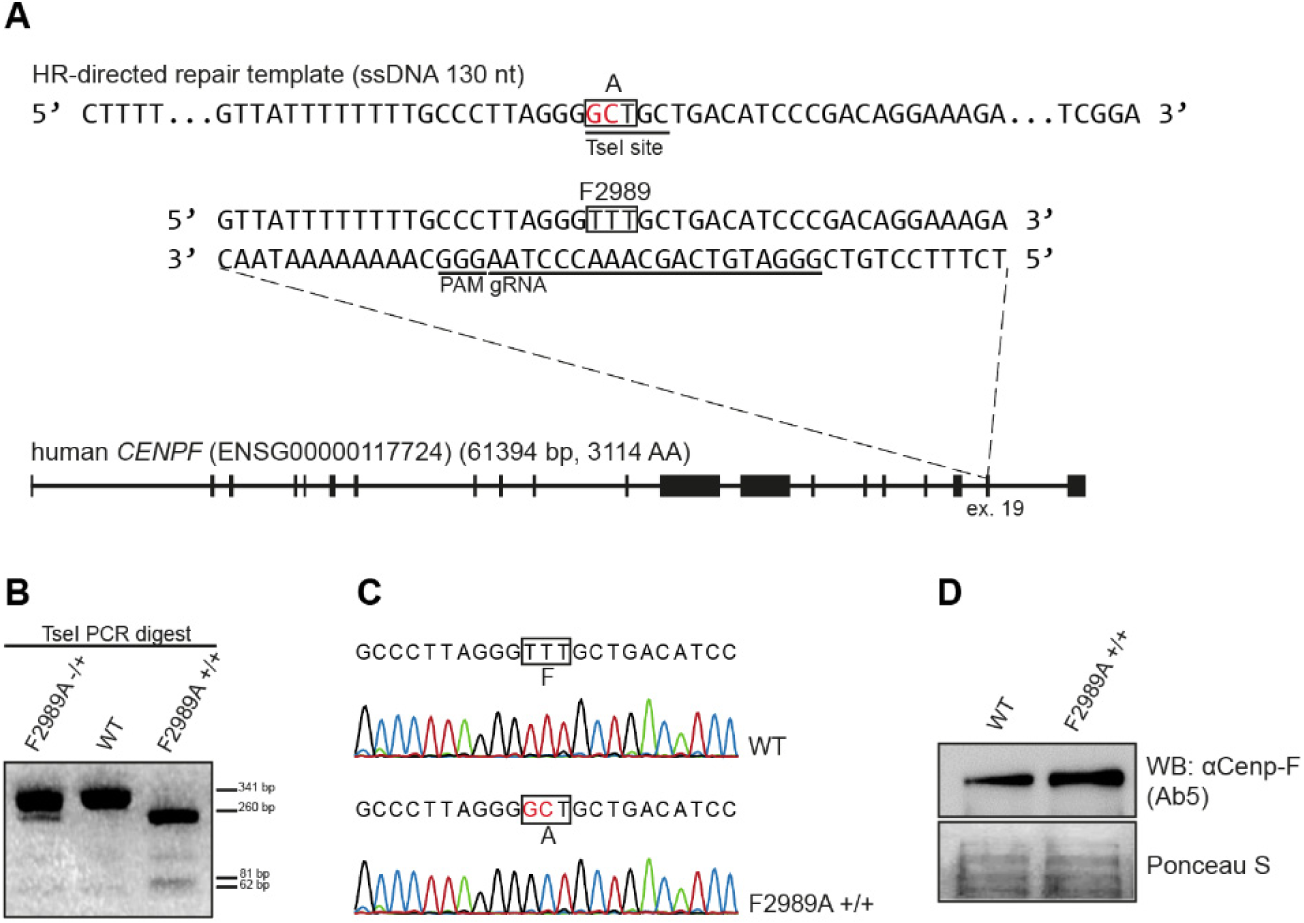
Introduction of the F2989A CENP-F mutation in human cells (**A**) Targeting strategy to engineer F2989A mutation in CENP-F using CRISPR/Cas9 in U2OS cells. 2nt substitution includes TseI restriction site to facilitate screening for correctly modified clones. (**B**) TseI digested PCR products of the targeted CENP-F locus from WT, heterozygous and homozygous CENP-F^F2989A^ U2OS cells. The expected restriction fragment sizes are 341 and 62 bp for the wild type allele, and 260, 81 and 62 bp for the F2989A allele. (**C**) Sequencing electropherograms of the targeted CENP-F area from WT and homozygous CENP-FF2989A U2OS cells. (**D**) Western blot on WT or CENP-F^F2989A^ U2OS cells using Ab5 antibody.

Our approach using co-electroporation of Cas9 RNP complexes and a 130nt single-stranded DNA homology-directed repair template yielded homozygous F2989A substitution in one out of 43 clones screened. 14 additional clones were heterozygous. This relatively high ratio of heterozygous versus homozygous modifications might be a consequence of polyploidy of U2OS cells (Janssen and Medema, 2012). To obtain additional homozygous clones for robust analysis, we re-targeted *CENP-F* in one heterozygous clone obtained in the first round of targeting. In this second round, 8 out of 26 clones were homozygous for CENP-F^F2989A^ (clones cl.2 to cl.9).

CENP-F^F2989A^ recapitulated the cell cycle-dependent expression and localization patterns of the WT CENP-F. It was well expressed (Figure 2D), accumulated in the nucleus during the S/G2 (Figure 3A), localized to the nuclear envelope in early prophase and to the kinetochores in mitosis (Figure 3B, C). However, while WT CENP-F could be seen accumulated on mitochondria in S/G2 and cytokinetic cells as previously described, CENP-F^F2989A^ remained diffuse in the cytosol throughout these cell cycle phases (Figure 3A). Taken together, these results indicate that F2989A mutation in CENP-F solely affects the mitochondrial localization of the protein.

**Figure 3:**
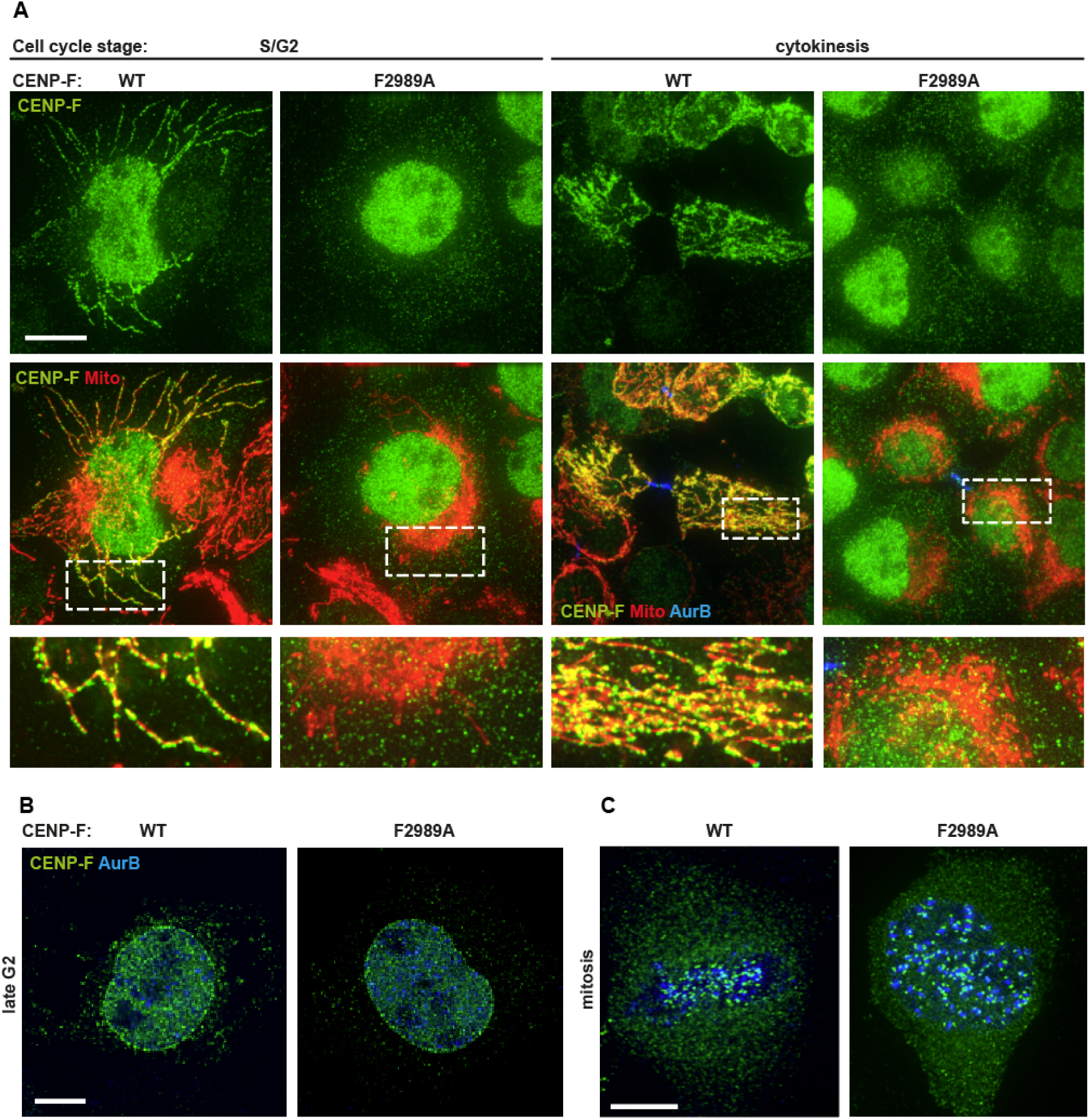
F2989A mutation disrupts CENP-F localization at mitochondria in human cells without affecting its localization at the nuclear envelope and kinetochores (**A**) Immunofluorescence of mitochondrial marker expressing (mtBFP) S/G2 or cytokinetic CENP-F^WT^ or CENP-F^F2989A^ cells using a CENP-F antibody (Ab5) (**B**) Immunofluorescence of CENP-F^WT^ or CENP-F^F2989A^ U2OS cells in late G2 using CENP-F (Ab5) and AuroraB antibodies. (**C**) Immunofluorescence CENP-F^WT^ or CENP-F^F2989A^ U2OS cells in mitosis using CENP-F (Ab5) and AuroraB antibodies. Scale bars 5 μm.

We have previously shown that siRNA-mediated depletion of CENP-F leads to defective distribution of the mitochondrial network throughout the cytoplasm. This defect most likely originates in cytokinesis during which CENP-F-depleted cells fail to redistribute mitochondria to the cell periphery (Kanfer et al., 2015). To validate that this defect was due to the absence of the mitochondrial pool of CENP-F, and not a side effect of global CENP-F depletion, we imaged mitochondria in the CENP-F^F2989A^ background. CENP-F^F2989A^ cells recapitulated the mitochondrial phenotypes of global CENP-F or Miro depletion in cytokinesis, displaying typical “clumped” mitochondria that failed to extend to the cell periphery (Figure 4A). Moreover, in interphase cells, mutant cells often displayed similar phenotype. This was not due to a failure of cell spreading, as bright-field imaging did not reveal aberrant cell shapes (Figure 4B). To quantify mitochondrial clumping in CENP-F^F2989A^ cells, we used an informatics approach that we have developed previously and that computes the “moment of inertia” of the mitochondrial network. This analysis measures the distance of each mitochondrial pixel to the cell’s center of mass, and thus, reflects the spreading of the mitochondrial network (Kanfer et al., 2015). Two independent clones (cl.1 and cl.2) bearing the homozygous CENP-F^F2989A^ mutation displayed a reduction in mitochondrial spreading, similar to what has been observed upon complete depletion of Miro or CENP-F (Figure 4C, and (Kanfer et al., 2015)). Therefore, the Miro-recruited mitochondrial fraction of CENP-F ensures that mitochondria are properly distributed throughout the cytoplasm in cytokinesis.

**Figure 4:**
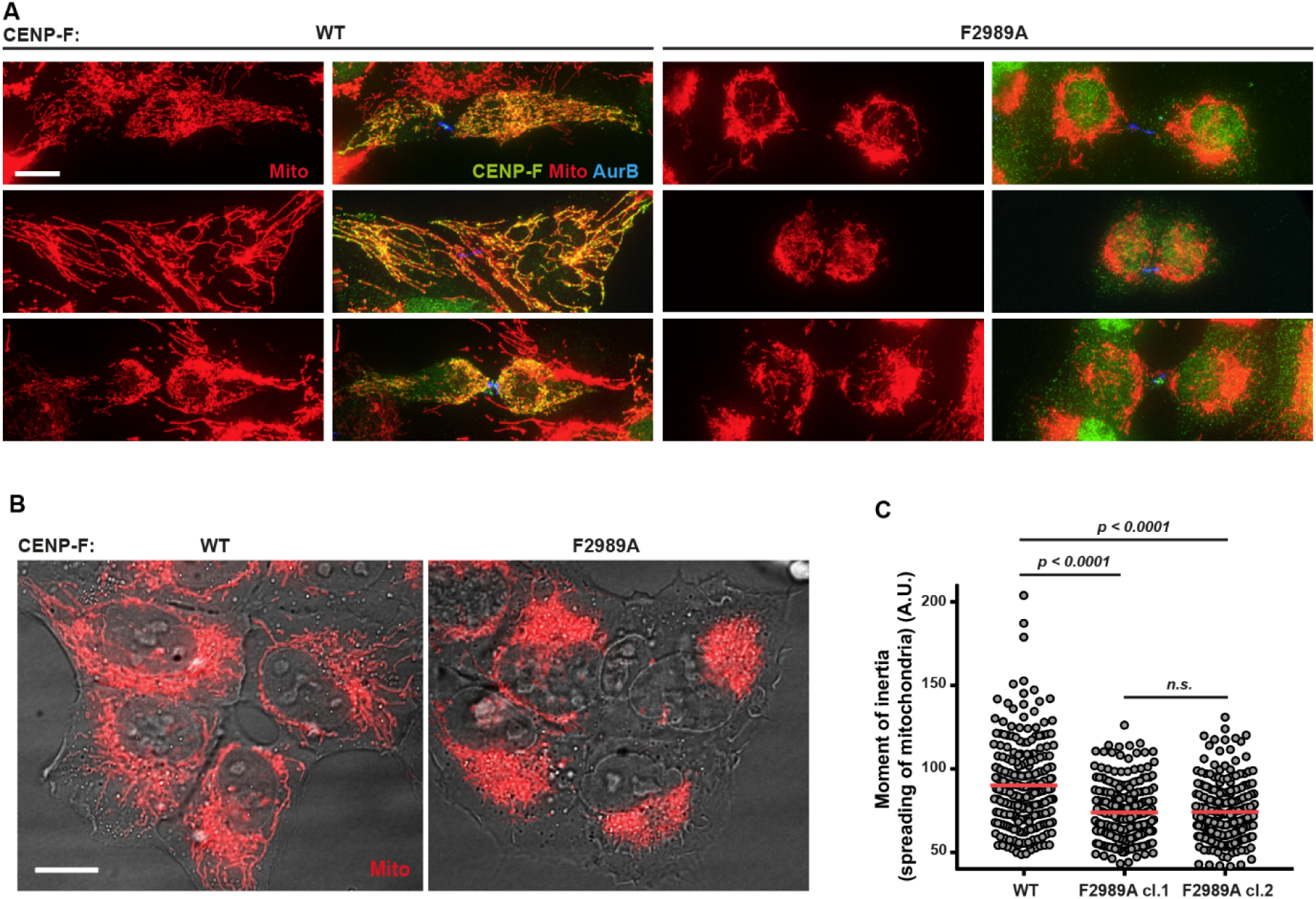
CENP-F-Miro complex is necessary for normal mitochondrial spreading in human cells (**A**) Immunofluorescence of mtBFP-expressing cytokinetic CENP-F^WT^ or CENP-F^F2989A^ U2OS cells using a CENP-F antibody (Ab5). (**B**) Examples of mitochondrial morphology in live CENP-F^WT^ or CENP-F^F2989A^ interphase U2OS cells stably expressing mtBFP. (**C**) Quantification of the moment of inertia (mitochondrial spreading) of cells exemplified in (B) Significance calculated using Mann-Whitney U test (N = at least 228 cells per condition) Two independent CENP-F^F2989A^ U2OS clones were used for quantification. Scale bars 5 μm.

To gain insight into the physiological function of mitochondrial CENP-F at the organismal level, we turned to genome editing in mice to mutate the conserved F2872 residue (cognate to human F2989) to alanine. For this purpose, we used pronuclear microinjection of preformed Cas9-gRNA complexes together with a single-stranded 130 bp homology-directed repair template (Figure 5A). In two microinjection sessions, we obtained 4 (20%) founders bearing heterozygous modification of the targeted F2872. These mutations were heritable, as crossing two heterozygous founders resulted in homozygous F1 mutant pups in expected Mendelian ratios (Figure 5B, Table 1). These homozygous animals (Figure 5C) were viable, fertile and reached adulthood without any apparent phenotype.

**Figure 5:**
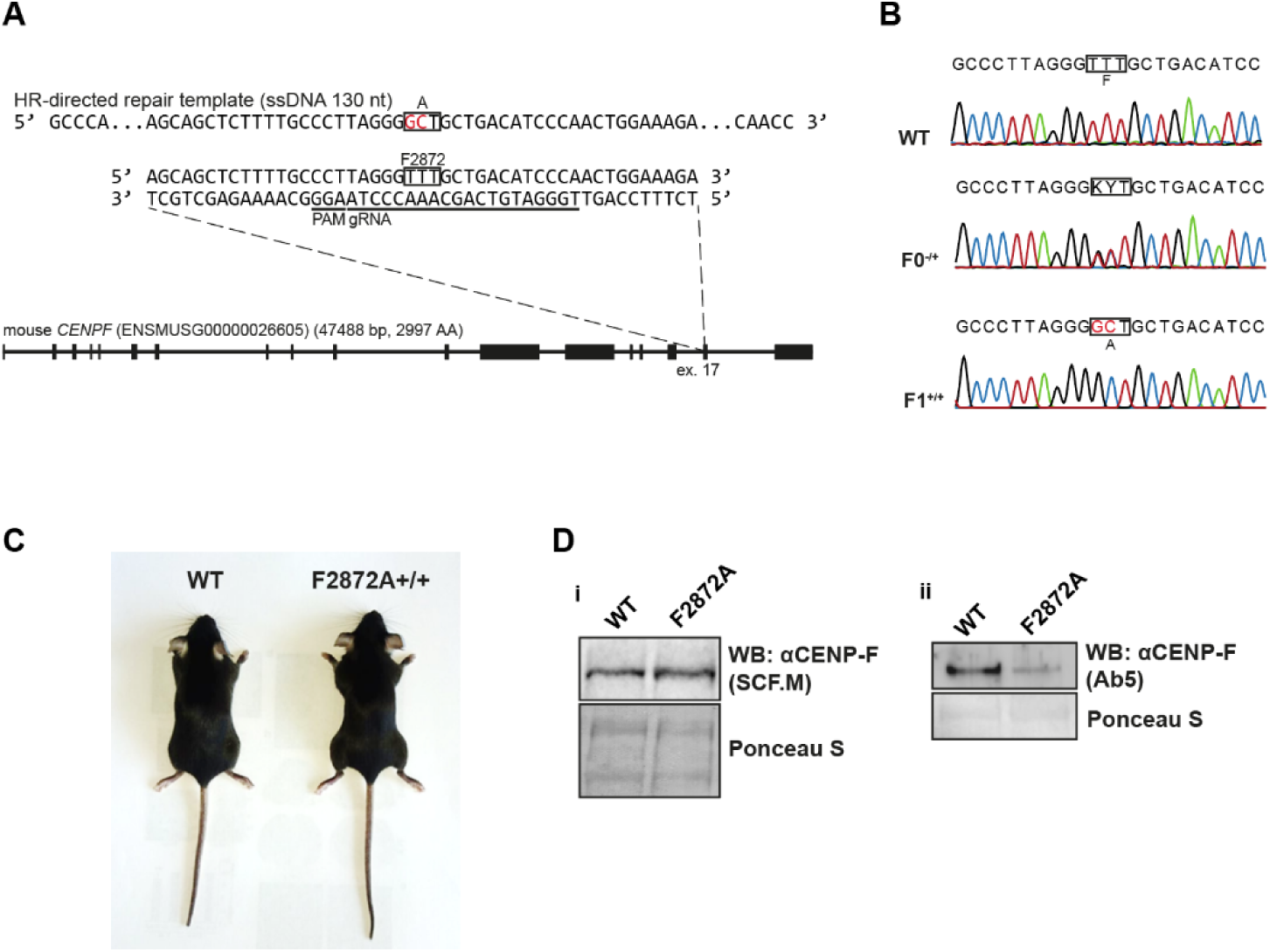
Introduction of the F2872A CENP-F mutation in mice (**A**) Targeting strategy to generate mice carrying F2872A in CENP-F using CRISPR/Cas9. (**B**) Sequencing electropherograms of the targeted CENP-F area from WT and F2872A-carrying F0 heterozygous or F1 homozygous mice. (**C**) 8 weeks old WT and homozygous F2872A litter-mate mice (**D**) Western blot on primary fibro-blasts derived from wild type or F2872A mice using (**i**) SCF.M or (**ii**) Ab5 antibody.

**Table 1:**
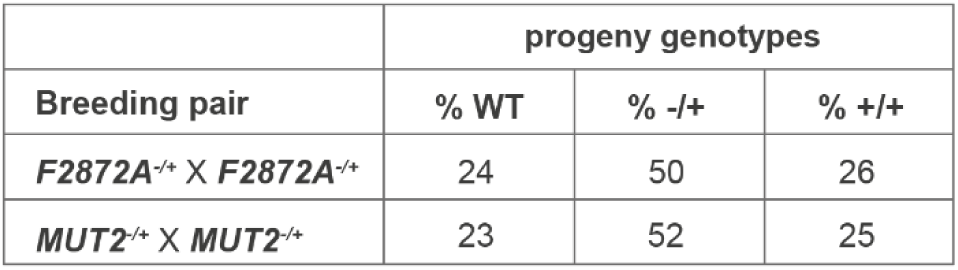
Heritability of genome-edited CENP-F alleles

We used western blotting with two different antibodies to examine if the expression level of CENP-F was affected by the F2872A mutation in fibroblasts derived from mutant animals. Using a polyclonal antibody specifically raised against the central area of mouse CENP-F (SCF.M antibody targeting residues 1408-1836. Courtesy of Stephen S. Taylor), we observed comparable expression levels between WT and the F2872A mutant (Figure 5Di). However, contrary to what we observed in the case of the mutant human cells (Figure 2D), we observed a reduced signal for the F2872A point mutant in western blot when using the widely used commercial CENP-F antibody Ab5 (Figure 5Dii). Ab5 is a polyclonal antibody raised against the C-terminal portion of human CENP-F (residues 2760-3114), an area that contains the mutated Miro-binding domain, suggesting that the F2872A mutation disrupts an important epitope.

Although raised against the human protein, Ab5 cross-reacts with the mouse protein, owing to the good primary sequence conservation in that area. In fact, the Miro-binding domain is the most conserved region between human and mouse species. The discrepancy between the SCF.M and Ab5 antibodies can therefore be explained as follows: the Ab5 polyclonal antibody preparation might contain antibodies recognizing several epitopes on the human protein, but only a limited set of epitopes on the mouse protein, among which, the Miro-binding domain is a preeminent one. Therefore, we conclude that the F2872A mutation does not affect protein stability but reduces the affinity of the Ab5 antibody for mouse CENP-F.

Since the presence of CENP-F on mitochondria has not been previously demonstrated in mice, we sought to validate the localization of CENP-F in cytokinetic mouse cells using immunofluorescence. The Ab5 antibody that we currently use for CENP-F immunofluorescence in human cells failed to generate a specific signal for murine CENP-F, consistent with a difference in epitopes between mouse and human proteins. We therefore performed all immunofluorescence experiments using the SCF.M antibody. Surprisingly, we did not detect any enrichment of WT CENP-F at mitochondria during cytokinesis in primary mouse fibroblasts (Figure 6A), but another commonly used mouse cellline NMuMG (mammary gland epithelium), displayed characteristic CENP-F enrichment on mitochondria during cytokinesis (Figure 6B). These data suggest that CENP-F-Miro interaction is cell-type specific and prompted us to examine CENP-F localization in primary epithelial cells derived from mammary gland. We thus harvested mammary glands from WT and mutant mice and extracted primary Mouse Mammary Epithelial Cells (MMECs) by digesting the tissue and culturing epithelial mammary gland organoids (Karantza-Wadsworth and White, 2008; McCaffrey and Macara, 2009). Immunofluorescence staining of CENP-F in these cells revealed strong mitochondrial enrichment of CENP-F during cytokinesis in WT cells. On the other hand, no such enrichment was observed in cells derived from CENP-F^F2872A^ mutant animals (Figure 6C), indicating that, as in human cells, the F2872 residue was essential for CENP-F-Miro interaction in mice. Nevertheless, unlike what was observed in U2OS cells, mitochondria spread equally well to the cell periphery, when compared to WT cells (Figure 6C). This discrepancy might be a consequence of differences in morphology between MMECs and U2OS cells. Unlike U2OS cells, MMECs grow in clusters and remain rounded throughout, and after, cytokinesis. Thus, mitochondrial spreading might not require CENP-F in MMECs.

**Figure 6:**
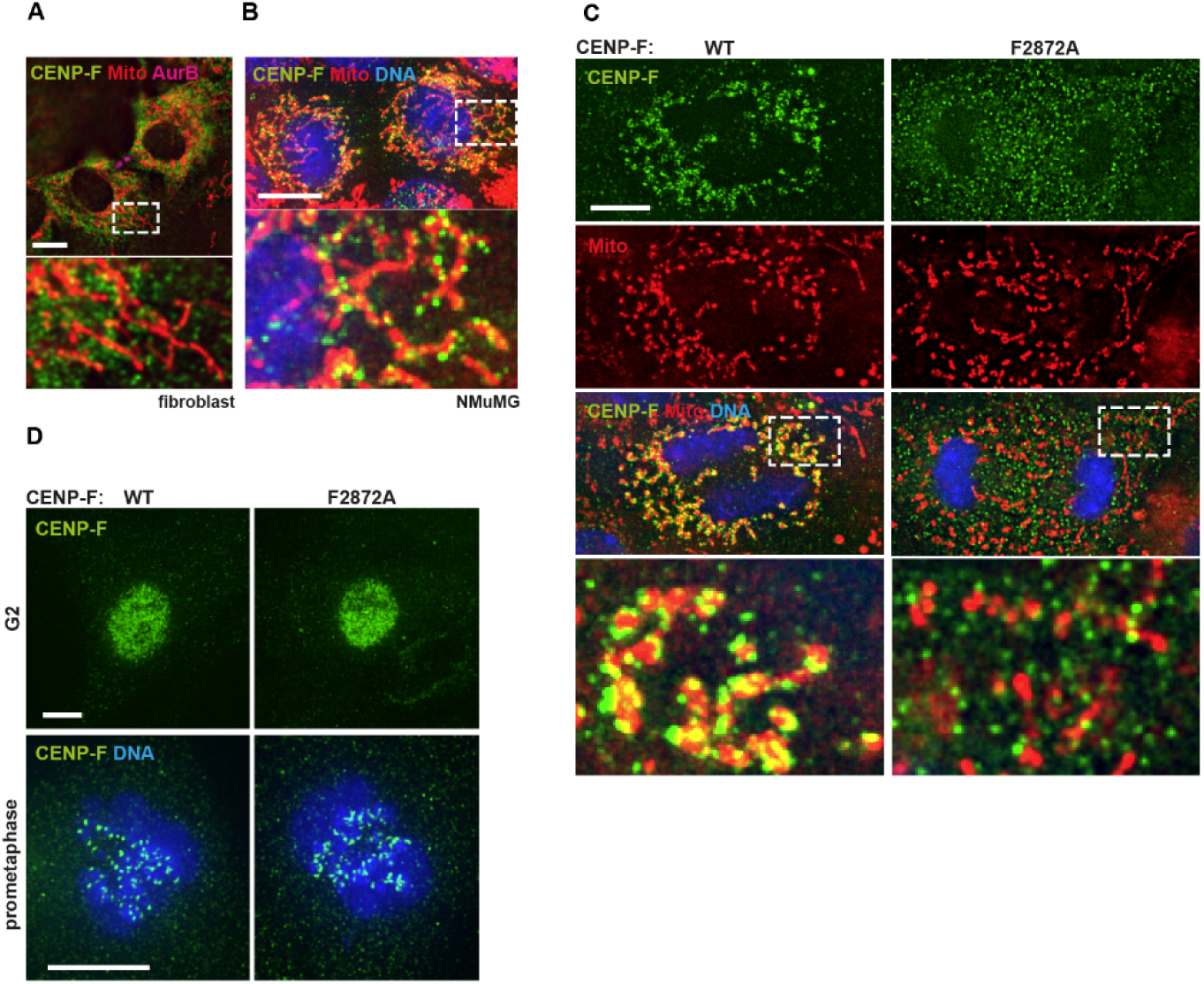
Mitochondrial CENP-F in mice is cell-type specific and can be disrupted by the F2872A mutation. (**A**) Immunofluorescence of mouse fibroblasts stained using CENP-F antibody (SCF.M), Aurora B antibody and mitotracker. (**B**) Immunofluorescence of mouse mammary epithelial cell line NMuMG stained using CENP-F antibody (SCF.M), mitotracker and DAPI. (**C**) Immunofluorescence of CENP-F (SCF.M antibody) in primary MMECs derived from WT or F2872A mice and labelled with Mitotracker and DAPI. (**D**) Immunofluorescence of CENP-F (SCF.M antibody) in primary MMECs derived from WT or F2872A mice and labelled with DAPI. Scale bars 5 μm.

Importantly, as in U2OS cells, CENP-F immunofluorescence revealed that CENP-F^F2872A^ is expressed and localized normally as shown by its nuclear accumulation in G2 and distinct kinetochore localization in mitosis (Figure 6D)

Taken together, these results demonstrate cell-type specific mitochondrial recruitment of CENP-F in mice, and show that CENP-F^F2872A^ is genetically uncoupled from mitochondria, but retains its other localizations.

As a consequence of homologous recombination being the limiting step in CRISPR knock-in experiments, we obtained two additional alleles that resulted from the repair of the Cas9-induced cuts by non-homologous end joining, in an error-prone manner (all mouse strains created here are summarized in Table 2). The first mutation was a 2bp deletion with a 4bp insertion at the very beginning of exon 17, which resulted in a frameshift causing a frameshift starting from residue G2871 (Table 2, Figure 7A, MUT1). The second allele contains a 13bp deletion encompassing the intron 16/exon 17 junction (Table 2, Figure 7A, MUT2). Thus, both of these mutants are expected to yield a truncation of the last two exons, which encode for (1) the C-terminal microtubule-binding domain of CENP-F, (2) part of the Miro-binding domain and (3) the CaaX farnesylation motif. The truncation may also affect the predicted Rb-interaction domain by removing its last five residues (Ashe et al., 2004). Both of these genotypes resulted in viable homozy-gous animals that are fertile and reached adulthood without any apparent phenotype (Figure 7B).

**Figure 7:**
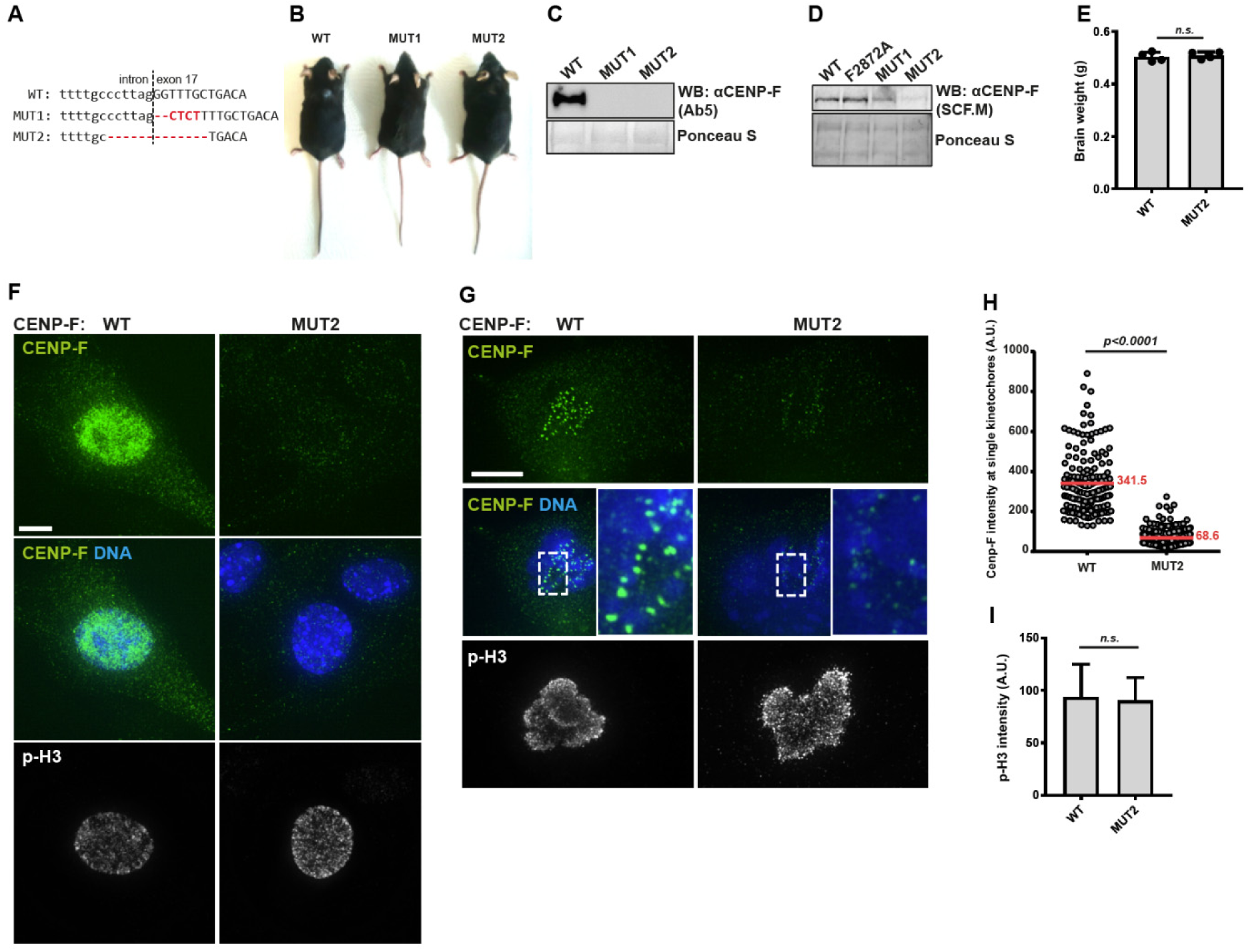
Engineering and characterization of CENP-F-truncating mutations in mice (**A**) CRISPR/Cas9-modified, indel-containing (red) CENP-F alleles in two different mice mutants (MUT1 and MUT2). (**B**) Adult WT and homozygous MUT1 and MUT2 mice. (**C**) Western blot using Ab5 antibody on primary fibroblasts derived from WT, MUT1 and MUT2 mice. (**D**) As in (C) but using the SCF.M. antibody. (**E**) Average brain weight of 8-week-old WT and MUT2 animals. Significance calculated using Mann-Whitney U test (N=4). (**F**) Immunofluorescence of late G2 primary fibroblasts from WT and MUT2 mice stained with CENP-F (SCF.M) and p-H3 S10 (06-570) antibodies, DNA labelled with DAPI. (**G**) Immunofluorescence of mitotic primary fibroblasts from wild type and MUT2 mice using CENP-F (SCF.M) and p-H3 S10 (06-570) antibodies, DNA labelled with DAPI. (**H**) Quantification of kinetochore CENP-F foci intensities in mitotic wild type or MUT2 cells exemplified in (E). At least 160 single kinetochores from 5 individual cells were quantified for each condition. Significance calculated using Mann-Whitney U test. Mean values depicted in red. (**I**) Mean intensities of p-H3 areas from mitotic cells exemplified in (E). Significance calculated using Mann-Whitney U test (N=5).

**Table 2:**
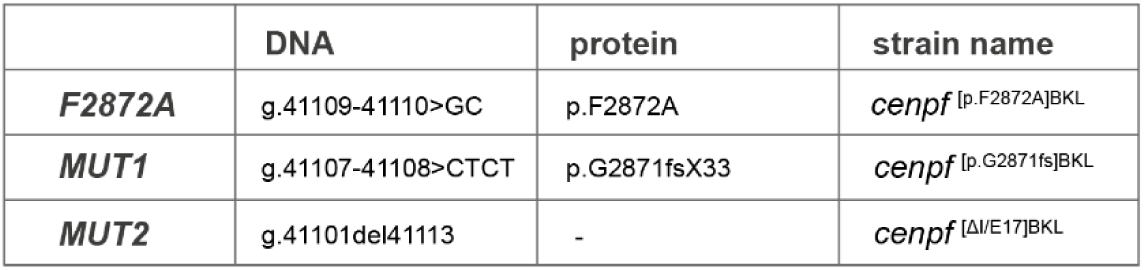
list of mouse lines generated in this study with description of modified alleles. Numbering of nucleotides and amino acids as in ENSMUSG00000026605

We first analyzed the expression levels of the mutant proteins using western blotting on primary fibroblasts. The Ab5 antibody raised against the C-terminus of CENP-F did not detect either allele, confirming that the C-terminus is missing in both mutants (Figure 7C). When using the SCF.M antibody raised against an epitope upstream of the mutations, we observed that both mutants showed reduced CENP-F levels. MUT2 displayed the most substantial loss of CENP-F expression (Figure 7D), prompting us to further analyze the abundance and localization of CENP-F throughout the cell cycle in these mutant animals.

We used Immunofluorescence staining to specifically observe the early mitotic population of cells, which were marked by histone H3 phosphorylation. While late G2/early M-phase WT cells displayed the expected nuclear accumulation of CENP-F, MUT2 cells did not display any CENP-F signal above background levels (Figure 7F). Nevertheless, we could detect residual CENP-F in prometaphase at kinetochores (Figure 7G). This allowed us to quantify the remaining fraction of MUT2 CENP-F by measuring fluorescence intensities of single CENP-F kinetochore foci. The mean fluorescence intensities (a.u.) of 341.5 in WT vs. 68.3 in MUT2 suggest an 80% loss of CENP-F expression in MUT2 (Figure 7H).

A recent report showed that CENP-F recruits ATR to kinetochores, which leads to activation of the Chk1-Aurora B axis, ultimately resulting in S10 phosphorylation of histone H3 and suppression of chromosome segregation errors (Kabeche et al., 2017). Our analysis did not show any decrease of pH3 S10 in MUT2-derived fibroblasts (Figure 7G, **I**).

CENP-F truncation in MUT2 is similar to alleles responsible for Strømme syndrome. A hallmark of Strømme syndrome patients is intestinal atresia, which becomes lethal unless resected. While we did not perform an in-depth phenotypic analysis of MUT2, the normal survival of these mice excludes the presence of intestinal atresia (Filges et al., 2016; Gao et al., 2009). Moreover, 8 weeks old MUT2 mice showed normal brain weight, suggesting no dramatic microcephaly phenotype, another characteristic of Strømme syndrome patients (Figure 7E).

## Discussion

CENP-F is an enigmatic protein with a plethora of suggested cellular functions ranging from chromosome segregation to mitochondrial trafficking. Because of the number of its binding partners and involvement in different cellular processes, a careful genetic dissection is needed to study specific roles of CENP-F. Variants of CENP-F have been characterized that separate its functions at the nuclear envelope and kinetochores (Berto et al., 2018; Zhu, 1999).

Here, we identify a point mutation (F2989A) in the conserved Miro-binding domain of CENP-F, which disrupts its interaction with Miro and thus its ability to be recruited to mitochondria. We thoroughly characterize the behavior of this variant by engineering it in cultured cells, and show that it is completely removed from mitochondria, while retaining other known localizations. To study the physiological relevance of the mitochondrial fraction of CENP-F, we mutated this residue also in mice and validated that CENP-F is removed from mitochondria. Surprisingly, these animals are viable and fertile with no apparent burden.

The transport of mitochondria is a crucial process in long cells like neurons. Its role in non-neuronal cells is much less clear. Preventing CENP-F-Miro interaction causes a drastic defect in mitochondrial spreading in U2OS cells, yet does not prevent the development of apparently healthy and fertile mice. This disconnect between *in vitro* and *in vivo* phenotypes suports a cell-type specific function of CENP-F-mediated mitochondrial trafficking. Indeed, we observe that mouse CENP-F is recruited to mitochondria in a Miro-dependent fashion in primary and immortalized mouse mammary epithelial cells, but not in primary mouse fibroblasts. This suggests that Miro-mediated CENP-F recruitment to mitochondria serves a tissue-specific rather than housekeeping function. What is this function? An in-depth phenotypic analysis of this mouse model is required to answer this question. Since the Miro-binding domain is one of the best conserved features of CENP-F, being almost identical from tunicates to human, it is likely that this function of the Miro-CENP-F interaction is also conserved. What is the molecular basis for the cell-type-specific Miro-CENP-F interaction? CENP-F Miro-binding domain is heavily phosphorylated and contains consensus sequences for several mitotic kinases. It is thus tempting to speculate that differential phosphorylation in different tissues and at different cell cycle stages promote or inhibit Miro-CENP-F interaction and the subsequent recruitment of CENP-F to the mitochondria.

While the mitochondrial function of CENP-F has been recognized only recently, many conflicting reports exist regarding its function in mitosis. Some studies advocate that CENP-F ensures correct chromosome segrega-tion by regulating kinetochore-microtubule attachments, thus affecting mitotic checkpoint and cell cycle progression (Bomont et al., 2005; Feng et al., 2006; Holt et al., 2005; Kabeche et al., 2017; Yang et al., 2005). On the contrary, other studies failed to detect any mitotic phenotypes upon CENP-F deletion in HeLa cells (McKinley and Cheeseman, 2017) or in mouse fibroblasts (Pfaltzgraff et al., 2016). Likewise, the function of the C-terminal farnesylation of CENP-F remains controversial. A number of reports showed diminished CENP-F localization at kinetochores upon farnesyltransferase inhibitor (FTI) treatment or upon mutagenesis of the CaaX motif (Gurden et al., 2010; Holland et al., 2015; Hussein and Taylor, 2002; Schafer-Hales et al., 2007). However, a recent study reported that FTI treatment or mutagenesis of the CaaX motif did not prevent kinetochore localization of CENP-F (Moudgil et al., 2015). Thus, further investigations into the function of CENP-F farnesylation are needed to clarify these conflicting observations.

Here, in addition to the F2872A mutant, we have generated mice carrying CENP-F alleles entirely lacking the C-terminal microtubule-binding domain and the farnesylation motif. Both of these have been implicated in possible CENP-F mitotic functions (Hussein and Taylor, 2002; Volkov et al., 2015). The normal survival of these animals excludes a drastic global defect in chromosome segregation, but it remains possible that certain cell types will be affected. Importantly, one of the mutated CENP-F alleles displays marked decrease of CENP-F expression. A recent report placed CENP-F as an important activator of mitotic checkpoint at the kinetochore, where it triggers a novel ATR - Chk1-AurB - pH3 pathway. The authors observed decrease in H3 phosphorylation upon expression of a presumed dominant-negative form of CENP-F. We did not see any decrease in H3 phosphorylation in fibroblasts derived from mutant animals. While it is possible that residual CENP-F in MUT2 might be fully capable of ATR activation, our observation is in line with the fact that CENP-F-null MEFs and HeLa cells do not display any defect in chromosome segregation (McKinley and Cheeseman, 2017; Pfaltzgraff et al., 2016), supporting that the mitotic function of CENP-F might be cell type or context specific.

Despite a host of important functions attributed to CENP-F, the protein appears mostly dispensable for development. The most straightforward explanation to this discrepancy lies in the fact that most studies to date have been performed in heavily transformed cancer cell lines. Some of these cell lines might have adapted their kinetochores to the fast pace of their divisions, possibly by becoming heavily dependent on CENP-F. Indeed, CENP-F emerges as a marker that is overexpressed in cancers (Aytes et al., 2014; Falagan et al., 2018; O’Brien et al., 2007; Zhuo et al., 2015). Here, we find that CENP-F can be downregulated without obvious consequences on animal health. Therefore, inhibiting CENP-F might be a viable avenue for treating CENP-F-dependent cancers.

CENP-F is also involved in human genetic diseases (Aytes et al., 2014; Filges et al., 2016; Ozkinay et al., 2017; Waters et al., 2015). In particular, familial mutations in human CENP-F lead to the Strømme syndrome, a disease characterized by severe ciliopathy phenotypes such as microcephaly and intestinal atresia. Interestingly, one of the non-functional CENP-F variants (p.Arg3094*) detected in Strømme patients is only predicted to lack the last 20 residues, indicating that minimal perturbations in CENP-F sequence can have drastic effects on human health (Filges et al., 2016). Some of the CENP-F mutant alleles found in Strømme patients were shown to be hypomorphic (Waters et al., 2015). We did not find a significant reduction in brain weight in young adult MUT2 animals. However, the possibility is open that more subtle changes in brain structure might be present. The mouse models presented here offer tools to study specifically the physiological functions of the mitochondrial fraction CENP-F, of the C-terminal microtubule-binding domain including the farnesylation site, and the consequences of global CENP-F downregulation. In the future, it will be important to examine the relevance of these mutants as potential models for CENP-F involvement in cancer and Strømme syndrome.

## Materials and methods

### Yeast two-hybrid assay

*LexA* operator-driven yeast two-hybrid assay was performed as described in (Golemis et al., 2011). Briefly, bait (pEG202-CENP-F^2977-3020^) and prey (pJG4-5-Miro1^1-594^) plasmids were constructed as described previously (Kanfer et al., 2015) and transformed into yeast strain EGY48. For the X-gal assay, transformants were plated on Gal/SD-His-Ura-Trp medium and assayed for LacZ reporter induction using the X-gal overlay method (Serebriiskii and Golemis, 2000). For the growth assay, transformants were plated on Gal/SD-Leu medium and grown for 48hrs at 30°C. CENP-F alanine point mutants G2988A, F2989A and T2997A were prepared by PCR-based site-directed mutagenesis of the CENP-F bait plasmid using primer pairs 1+2, 3+4 and 5+6 respectively (all primers listed in Table 3).

**Table 3:**
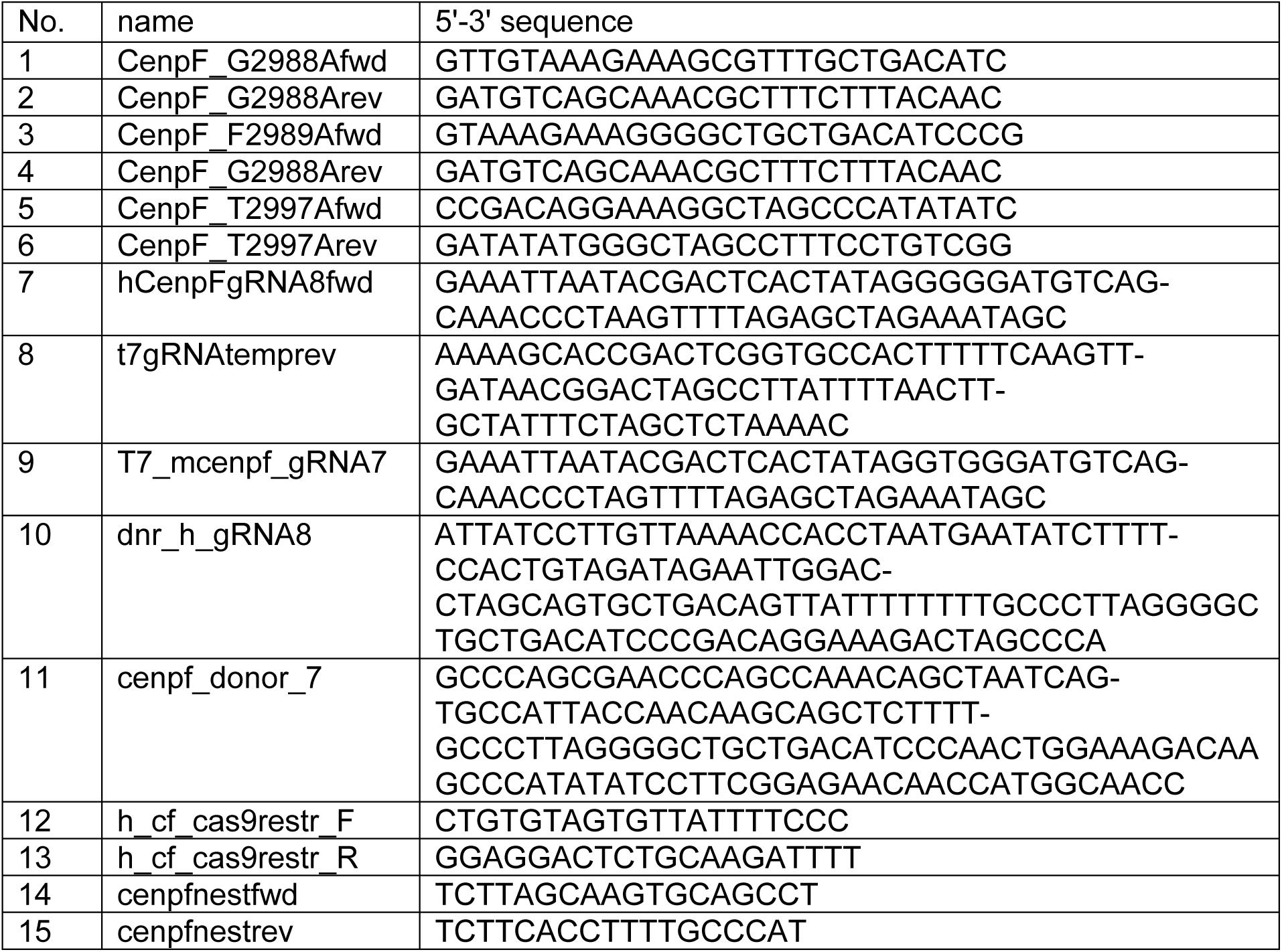
list of oligonucleotides used in this study

### Immunofluorescence

For immunofluorescence staining (IF) using the αCENP-F (Ab5, Abcam), αAuroraB (Clone 6/AIM-1, BD Biosciences) and p-H3 (S10) (Milipore, 06-570) antibodies, cells were grown on coverslips to reach ∼80 % confluency and fixed using PBS/4 % PFA for 30 minutes at room temperature. Subsequently, the samples were washed three times with PBS/0.1 % Triton-X100 and incubated in the blocking solution (PBS/10 % FCS) for 30 minutes. The samples were then incubated with primary antibodies for 1 hour (dilution 1:300 in blocking solution), washed three times with PBS/0.1 % Triton-X100 and incubated with fluorophore-conjugated secondary antibodies (dilution 1:1000 in the blocking solution) for 30 minutes. Finally, the samples were washed three times with PBS/0.1 % Triton-X100, three times with PBS and then mounted to slides using Vectashield mounting medium, optionally containing DAPI to visualize DNA. For immunofluorescence staining of mouse cells using SCF.M antibody (kind gift from Stephen Taylor, The University of Manchester), the same steps were performed except that cells were fixed with ice-cold methanol (-20°C, 10 minutes). To visualize mitochondria, MitoTracker Deep Red FM (Invitrogen/Molecular Probes) was added to the growth media (100nM, 45 minutes) before cell fixation. Imaging was performed using a DeltaVision wide-field microscope (Olympus IX71) equipped with 100x 1.4NA Oil UPlanSApo objective, DAPI-FITC-TRITC-CY5 filter set (Chroma) and Roper CoolSnap HQ2 camera. Images were deconvoluted with Softworx 4.1.0 (Applied Precision). Final image processing and analysis were performed using Fiji (Schindelin et al., 2012).

### Quantification of mitochondrial spreading

For quantification of mitochondrial spreading, live U2OS cells stably expressing mtBFP (Kanfer et al., 2015) were grown and imaged in Labtek chambers. Images of mitochondria were acquired at 37°C using the wide-field microscope setup described above. For each image of the mitochondrial network, the moment of inertia (equivalent of mitochondrial spreading) was calculated using custom Fiji and Matlab scripts as detailed previously (Kanfer et al., 2015).

### Quantification of CENP-F kinetochore localization and p-H3 signal intensity

To quantify residual CENP-F at kinetochores, CENP-F MUT2 and WT mitotic fibroblasts were fixed and labelled using the SCF.M αCENP-F antibody, and DAPI to visualize mitotic chromosomes. Z-sections of prometaphase cells were acquired in 0.3 µM steps using the wide-field microscope described above and then deconvoluted. Images were subsequently processed and analyzed in Fiji. In short, the sections were Z-projected (maximum intensity) and kinetochore CENP-F signal was calculated using the *IntDen - (Area * Mean)* formula (*IntDen* = integrated signal density of a single kinetochore, *Area* = the size of the selected kinetochore area and *Mean* = the mean fluorescence of the CENP-F signal outside the kinetochores). At least 164 single kinetochores were measured for each condition. To quantify phosphorylation of histone H3 phosphorylation in mitotic cells, MUT2 and WT fibroblasts were immunostained for p-H3 (S10) and processed similarly. Mean fluorescence intensities of p-H3 (S10) areas in mitotic cells were measured and normalized to the background.

### Western blotting

Primary skin fibroblasts or U2OS cells were harvested and lysed in 8M urea with 3 % SDS. Equal amounts of protein lysates as measured by the Bradford assay (Bio-Rad) were resolved by SDS-PAGE and analyzed by western blotting using Ab5 or SCF.M antibodies (1:500 dilution

### Generation of CRISPR/Cas9-edited U2OS cells

*CENP-F* was targeted using electroporation (Neon system, Invitrogen) of Cas9 RNPs and DNA template for homologous recombination (also see Figure 2 for strategy details). Briefly, ∼200.000 cells were harvested, washed with PBS and resuspended in 5 µl of Neon R buffer (Invitrogen). Shortly before electroporation, Cas9 RNP mix (100 pmol of Cas9 (NLS-Cas9, a kind gift from Martin Jinek, University of Zurich), 120 pmol of sgRNA and 200 pmol of single stranded donor DNA template (10, Table 3) in total volume of 5 µl of Neon R buffer) was added to the cell suspension and electroporation was performed using 10 µl Neon tips with electroporation parameters set to 1400V, 15 ms, and 4 pulses. Immediately after electroporation, cells were seeded into 0.5 ml of pre-warmed growth medium and grown for 3 days. Cells were then single-cloned and colonies arising from single clones were used for PCR amplification of the targeted *CENP-F* locus using primers 12+13. The resulting PCR product was sequenced using primer 12 or digested by TseI to screen for the intended modification (Figure 2B). Clones containing homozygous F2989A mutations in *CENP-F* were collected for further analysis.

### Generation of CRISPR/Cas9-edited CENP-F mutant mice

Mice were generated and all animal work was performed in the animal facility of ETH Zurich (EPIC) according to the Swiss Federal Veterinary Office (BVET) guidelines. Animals were regularly checked for potential burden. *CENP-F* knock-in and truncated alleles in mice (B6D2F1 x C57Bl/6NTac mixed background) were generated using a modified Cas9-RNPs approach (Sung et al., 2014). gRNA sequence targeting the proximity of the CENP-F F2872 codon was selected using the CRISPR design tool (http://crispr.mit.edu/) (also see Figure 5A for details of the targeting strategy). *In vitro* transcribed sgRNA was prepared as described below. ssDNA containing F2872A codon change flanked by 60nt homology arms was used as a template for homologous recombination (11, Table 3). Injection mix was assembled on ice by mixing 200 nM of Cas9-NLS (NEB), 400 nM sgRNA and 100 ng/µl ssDNA donor in the injection buffer (8 mM Tris-HCl, pH 7.4, 0.1 mM EDTA). The mix was injected into pronuclei of fertilized single-cell embryos using standard microinjection procedures. Injected embryos were re-implanted into foster mothers. The pups were genotyped by PCR on genomic DNA extracted from ear clips using primers 14+15. The resulting PCR product was sequenced using primer 14.

### *In vitro* transcription of sgRNAs

sgRNAs were prepared using MEGAshortscript T7 Transcription Kit and MEGAclear Transcription Clean-Up Kit (Ambion) according to manufacturer instructions. The DNA template for *in vitro* transcription was generated by PCR-assembly of partially overlapping oligonucleotides containing T7 promoter sequence, mouse or human CENP-F gRNA target sequence (Figure 2A and 5A) and an invariant Cas9 scaffold sequence (human: primers 7+8, mouse: primers 9+8).

### Primary mouse cells isolation

Primary skin fibroblasts were isolated as described step-by-step in (Seluanov et al., 2010). In brief, approximately 1cm^2^of skin was collected from the underarm area of freshly euthanized ∼ 8-week-old animals, cut into small pieces and dissociated in 10ml of Liberase TL (Roche) solution (0.14 Wunsch units/mL in DMEM/F12 + 1X antibiotic/antimycotic) at 37°C for 1 hour while stirring. Dissociated tissue fragments were collected by centrifugation (520g/5 minutes) and washed three times in DMEM/F12 with 15 % FBS to remove remaining Liberase. The tissue fragments were then plated in DMEM/F12 with 15 % FBS + 1X antibiotic/antimycotic and grown at 37°C, 5 %CO_2_ for 7 days allowing fibroblasts to exit the tissue. Mouse mammary epithelial cells (MMECs) were isolated as described in (Karantza-Wadsworth and White, 2008; McCaffrey and Macara, 2009). In brief, 3^rd^, 4^th^ and 5^th^ pairs of mammary glands were collected from freshly euthanized ∼ 8-week-old animals, minced using scissors and dissociated in Collagenase A (Roche) solution (2 mg/ml in DMEM/F12 + 1X antibiotic/antimycotic) at 37°C for 2 hours while stirring. The dissociated epithelial organoids were pelleted from the suspension by centrifugation (250g/5 minutes) and resuspended in 5ml DMEM/F12. The organoids were again pelleted (250g/5 minutes) and resuspended in 10ml PBS/5 % FBS. The pellet was then resuspended and washed with 10ml PBS/5 % FBS five times (400g/15 seconds) to remove single cells from the organoids. After the last washing step, the pellet containing pure epithelial organoids was resuspended in DMEM/F12, 10 % FBS, 5 µg/ml insulin (Sigma, I0516), 1 µg/ml hydrocortisone (Sigma, H0888), 5 ng/ml hEGF (Sigma, E9644) + 1X PSG mix) for 48 hours and switched to serum-reduced medium (DMEM/F12, 5 % FBS, 5 µg/ml insulin, 1 µg/ml hydrocortisone, 5 ng/ml EGF + 1X PSG mix) to prevent growth of remaining fibroblasts. Epithelial cells escaped the organoids within 2-3 days and were immediately processed for immunofluorescent staining.

## Acknowledgments

We are grateful to the EPIC facility of ETH Zurich& Thomas Hennek for help with generating and handling CRISPR-edited mice, Martin Jinek (University of Zurich) for kindly providing recombinant Cas9 and Stephen S. Taylor (The University of Manchester) for kindly sharing the SCF.M antibody. We also thank all members of the Kornmann lab for providing comments on the manuscript. Microscopy was performed at the Scientific Center for Optical and Electron Microscopy of the ETH Zurich. This work was funded by grants from the ERC (337906-Orga-Net) and the Swiss National Science Foundation (PP00P3_13365) to BK.

